# Alterations in cortical thickness and structural connectivity are associated with symptom severity in bulimia nervosa

**DOI:** 10.1101/127910

**Authors:** Margaret L. Westwater, Jakob Seidlitz, Kelly M.J. Diederen, Sarah Fischer, James C. Thompson

**Affiliations:** Department of Psychiatry, University of Cambridge, Herchel Smith Building, Addenbrooke’s Hospital, Cambridge CB2 0SZ, UK; Department of Psychology, George Mason University, Fairfax, VA 22030 USA

**Keywords:** Eating disorders, structural MRI, symptom severity, frontoparietal control network, orbitofrontal cortex

## Abstract

Bulimia nervosa (BN) is a serious psychiatric illness defined by preoccupation with weight and shape, episodic binge-eating and compensatory behaviors. Although diagnosed BN has been associated with diffuse grey matter volume reductions, characterization of brain structure alterations in women with a range of BN symptoms has yet to be made. This study examined whether changes in cortical thickness (CT) scaled with BN symptom severity in a sample of 33 adult women (n = 10 BN; n = 5 EDNOS-BN). Our second objective was to assess global structural connectivity (SC) of CT and to determine if individual differences in global SC relate to BN symptom severity. We used the validated Eating Disorder Examination Questionnaire (EDE-Q; Fairburn & Beglin, 1994) as a continuous measure of BN symptom severity. Increased EDE-Q score was negatively related to global CT and local CT in the left middle frontal gyrus, right superior frontal gyrus and bilateral orbitofrontal cortex (OFC) and temporoparietal regions. Moreover, analysis of global SC indicated that BN-related cortical thinning preferentially occurred in regions with high global connectivity. Finally, we showed that individuals’ contribution to global SC at the group level were significantly related to EDE-Q score, where increased EDE-Q score correlated with reduced connectivity of the left OFC and middle temporal cortex and increased connectivity of the right superior parietal lobule. Our findings offer novel insight into CT alterations in BN and further suggest that the combination of CT and structural connectivity measures may be sensitive to individual differences in BN symptom severity.

## 1. Introduction

Characterized by recurrent binge-eating, compensatory behaviors to preclude weight gain, and over-valuation of body weight and shape, bulimia nervosa (BN) is a serious psychiatric disorder that afflicts 1 − 2.3% of the population [Hoek and van Hoeken, 2003; Keski-Rahkonen et al., 2009]. BN typically arises in adolescence or emerging adulthood [Fairburn and Harrison, 2003; Favaro et al., 2009], and affected individuals demonstrate altered brain activity in response to self-regulatory (e.g., Go/NoGo, Stroop; Marsh et al., 2009), gustatory [Bohon and Stice, 2011] and body processing [Vocks et al., 2010] tasks. Research on normative adolescent development has largely characterized maturation of the cortex, where rapid cortical changes are thought to underpin the increases in cognitive and behavioral capacity that occur throughout adolescence [Knudsen, 2004]. However, this critical period may also increase risk for psychiatric disease [Giedd, 2008]. The average age of onset of BN, paired with reported alterations in functional brain activity, could imply changes in underlying brain structure.

Although several studies have examined associations between BN and brain structure, particularly grey matter volume, findings are inconclusive. For instance, some voxel-based morphometry (VBM) studies of women with BN report increased orbitofrontal and ventral striatal grey matter volume [Frank et al., 2013; Schäfer et al., 2010] when compared to healthy control women. However, manual tracing of striatal structures suggests volumetric reduction of the caudate nucleus and preservation of nucleus accumbens volume among women with BN [Coutinho et al., 2015]. Examination of group differences across adolescent and adult women has shown reduced local volumes in frontal and temporoparietal regions in bulimic individuals [Marsh et al., 2015]. The authors also reported diffuse cortical thinning in the BN group, where the most significant reductions were seen in bilateral precentral and inferior frontal gyri. When exploring the relationship between symptom severity and cortical volume, Marsh et al. [2015] reported a negative association between bingeing and vomiting frequency and volume of inferior frontal and central gyri. Exploratory analyses further indicated reduced grey matter integrity in inferior, middle frontal and central gyri in women with BN who endorsed heightened shape and weight concerns. Interestingly, none of these studies observed differences in whole-brain [Marsh et al., 2015) and total grey matter [Frank et al., 2013; Wagner et al., 2006] volume in individuals with BN.

Such inconsistencies in regional grey matter alterations may arise, in part, from the use of different analytic techniques. While volumetric analysis can be informative when examining deep brain and subcortical structures, the interpretability of volumetric changes in the cortex is limited, as this composite measure comprises two features—thickness and surface area—that arise from distinct genetic factors [Panizzon et al., 2009; Winkler et al., 2010]. Furthermore, replication studies indicate that surface area, not cortical thickness, explains a greater proportion of grey matter volume variance in adults [Im et al., 2008; Pakkenberg and Gundersen, 1997]. These and other methodological challenges may be overcome by surface-based morphometric analysis, which delineates the distinct features of the cortex, enabling a more precise interpretation of underlying cytoarchitecture [Wagstyl et al., 2015].

In addition to potential limitations of volumetric morphometry analysis, the use of diagnostic cut-offs and case-control designs may not fully characterize dysfunctional neural circuitry in BN. Psychiatric nosology has begun to shift toward a dimensional approach [Insel et al., 2010], largely in response to research illustrating that psychiatric disorders exist on a continuum, are chronic, and often exhibit high comorbidity with one another [Gandal et al., 2016]. Thus, categorical approaches to psychiatry may not capture the full range of individual differences between affected individuals, which could result in the exclusion of meaningful variance in brain measures. Variability in brain measures may contribute to heterogeneity in the clinical presentation of BN, and the use of a dimensional approach, which examines the full distribution of a given trait or characteristic, can therefore aid the translation of neuroimaging research to clinical practice. Several lines of evidence support a dimensional model of bulimic pathology [Stice et al., 1996; Stice et al., 1998], where continuity of cognitive, affective and behavioral components has generally been evaluated through group comparisons of healthy, ‘subclinical’ bulimic, and bulimic women. Although results from preliminary functional magnetic resonance imaging (fMRI) studies of reward processing support a dimensional model in women with full and subclinical BN [Bohon and Stice, 2011], the question of whether anatomical alterations scale with symptom severity in BN remains unanswered.

The present study utilizes a dimensional approach to assess alterations in cortical thickness (CT) across adult women with a wide range of BN pathology. We have selected CT as a reliable morphometric measurement, which can be measured at a high degree of spatial resolution across the entire cortical mantle [Fischl and Dale, 2000]. Moreover, both cross-sectional [Lerch et al., 2006; Whitaker et al., 2016] and prospective [Alexander-Bloch et al., 2013a; Raznahan et al., 2011] studies of normative development indicate that focal changes in CT occur in a coordinated manner across the cortex. This would suggest that regional alterations in CT are not statistically independent but are instead correlated with one another. As such, we also aimed to quantify the structural connectivity of CT across the cortex at the group level and examine how individual contributions to this connectivity [Saggar et al., 2015] are related to BN symptom severity. To provide a continuous measure of disordered thoughts about weight, shape and eating, BN symptoms were quantified using a validated self-report measure, the Eating Disorder Examination Questionnaire [EDE-Q; Fairburn and Beglin, 1994]. We hypothesized that BN symptom severity would be negatively related to both global and local measurements of cortical thickness, localized to frontal, temporal and parietal lobules.

## 2. Materials and Methods

### 2.1 Design and Participants

Thirty-seven adult women (M_age_ ± SD; 22.6 ± 4.13 y, 62% White) with a range of BN symptoms participated in the behavioral and scanning portions of the study. Participants were recruited from George Mason University, an outpatient eating disorder clinic and the local community via online and posted advertisements. Within the sample, 17 subjects completed a diagnostic phone interview as a part of a separate study, which was used to determine the presence of bulimic symptoms and compatibility for MRI scanning. An extended description of participant recruitment can be found in Fischer et al. [2017]. The remaining 20 subjects were initially screened for contraindications to MRI scanning and invited to participate in the behavioral and scan sessions. All participants were right-handed with normal or corrected-to-normal vision. Exclusion criteria included neurological disorders or traumatic brain injury, metallic implants, psychosis or substance dependence, and pregnancy. Participants provided written, informed consent in accordance with George Mason University Human Subjects Review Board guidelines and received monetary compensation for their time.

To determine the presence of BN symptoms, all subjects completed the Eating Disorder Examination [EDE; Cooper and Fairburn, 1987] with a trained clinician. The EDE is a semistructured clinical interview that provides an initial diagnosis of anorexia and bulimia nervosa, binge-eating disorder (BED) and eating disorder not otherwise specified (EDNOS) based on the Diagnostic and Statistical Manual of Mental Disorders 4th edition [DSM-IV; American Psychiatric Association, 2000] criteria. Subjects also completed screening questions from substance dependence and psychosis modules of a structured clinical interview for DSM-IV Axis I psychiatric disorders [First et al., 2002]. Following the interview, participants completed the EDE-Q to provide a continuous measure of disordered eating symptoms and behaviors. Scores for each EDE-Q subscale (eating concern, weight concern, shape concern and restraint) were computed and averaged to generate a mean symptom severity score (greater scores indicate increased severity). We operationalized BN symptom severity with EDE-Q score. Because we oversampled for women with BN symptoms and validated these symptoms with the EDE, EDE-Q score was thought to closely reflect severity of BN as opposed to any eating disorder. Self-report data related to depressive [Quick Inventory of Depression; Rush et al., 2003] and anxiety [State and Trait Anxiety Inventory; Spielberger and Sydeman, 1994] symptoms were also collected. These data and demographic information are presented in Table 1.

**Table 1.** Clinical and demographic information.

| Characteristic | Sample (N = 37) |  |
| --- | --- | --- |
|  | M ± SD | Range |
| Age, y | 22.6 ± 4.13 | 18 – 38 |
| BMI, kg/m<sup>2</sup> | 23.9 ± 3.14 | 19.4 – 31.2 |
| BN, # (%) | 12 (32.4) |  |
| EDNOS with BN symptoms, # (%) | 5 (13.5) |  |
| Medication, # (%) | 2 (5.4) |  |
| EDE-Q Total Score | 2.28 ± 1.53 | 0.0 – 5.04 |
| *Restraint* | 2.01 ± 1.48 | 0.0 – 4.40 |
| *Eating Concern* | 1.53 ± 1.55 | 0.0 – 5.0 |
| *Weight Concern* | 2.70 ± 1.87 | 0.0 – 5.80 |
| *Shape Concern* | 2.90 ± 1.76 | 0.0 – 5.75 |
| Objective Binge Episodes | 3.34 ± 4.91 | 0 – 20 |
| Subjective Binge Episodes | 2.13 ± 3.41 | 0 – 12 |
| Vomiting Episodes | 0.85 ± 2.52 | 0 – 12 |
| Laxative Use | 0.91 ± 2.73 | 0 – 14 |
| Excessive Exercise Episodes | 6.29 ± 8.97 | 0 – 28 |
| QIDS Score | 8.16 ± 5.56 | 1 – 21 |
| TAI Score | 46.3 ± 11.35 | 27 – 70 |

**Notes:** BMI = body mass index; BN = bulimia nervosa; EDNOS = eating disorder not otherwise specified; QIDS = Quick Inventory of Depressive Symptoms; TAI = Trait Anxiety Inventory

### 2.2 MRI Data Acquisition & FreeSurfer Reconstruction

MRI data were collected on a Siemens 3T Allegra scanner (Erlangen, Germany) with a one-channel, quadrature birdcage head coil. T1-weighted structural scans were obtained using a three-dimensional, magnetization-prepared, rapid-acquisition gradient echo (MPRAGE) pulse sequence with sagittal acquisition (160 1-mm thick slices, flip angle = 9°, matrix size = 256×256, FOV = 260 mm2, TR = 2,300 ms, TE = 3.37 ms). All scans were inspected for motion artifacts and reviewed by a neuroradiologist for gross anatomical abnormalities.

Using MRIcron [Rorden and Brett, 2000], DICOM images were converted to NIFTI files. To extract CT measurements, cortical surface reconstructions were generated in the FreeSurfer software package (v. 5.2; http://surfer.nmr.harvard.edu), following the procedures described by Dale et al. [1999] and Fischl et al. [1999]. In brief, MR images were corrected for magnetic field inhomogeneities, affine registered to the Talairach atlas [Talairach and Tournoux, 1988] and skull stripped to remove non-brain tissue. White matter (WM) voxels were masked using a six-neighbors connected components algorithm and separated by hemisphere for preliminary segmentation. A triangular mesh was then fitted to the WM surface, tessellated and smoothed with a Gaussian kernel to provide a realistic representation of the grey-white matter boundary. The WM surface was subsequently deformed outward to the point of maximal tissue contrast to determine the pial surface. Cortical thickness measurements were computed as the shortest mean distance between the WM and the pial surface, where thickness was measured first from the WM to the pial surface and again in reverse from the pial to the WM surface. Thickness values were generated for each vertex on the subject’s surface and averaged across each hemisphere to determine mean CT. Completed cortical surface reconstructions were visually inspected for quality control purposes, and manual editing of WM and non-brain tissue (e.g., dura mater) was performed by MLW where necessary.

### 2.3 Global & Local Cortical Thickness Analyses

Linear regression analysis of the association between BN symptom severity and global CT was completed in R [R Core Team, 2015]. We calculated global CT for each subject using the normalization technique described by Winkler and colleagues [2010]. This procedure first weights the mean thickness of each hemisphere by the corresponding surface area and then sums the values. Accounting for surface area effectively controls for inter-individual differences in brain size. The linear regression model examined the relationship between EDE-Q score and global mean CT, with age and BMI entered as covariates. Normality of the model residuals was determined by the Shapiro-Wilk test.

Vertex-wise analysis of local changes in CT related to BN symptom severity was performed using the general linear modeling (GLM) application in FreeSurfer. Using the GLM, thickness values were modeled as a linear combination of the effects of the predictor variable(s) at each vertex. EDE-Q score was entered as a continuous predictor in the model, and age (y), body mass index (BMI; kg/m^2^) and WM surface area (mm^2^) were entered as covariates of no interest. Each subject’s surface was registered to the fsaverage surface template and smoothed with a 15 mm FWHM Gaussian kernel. We examined a single GLM matrix in each hemisphere, which generated a t-statistic map of the correlation between EDE-Q score and thickness, controlling for age, BMI and brain size. To correct for multiple comparisons, a Monte Carlo Null-Z simulation (10,000 iterations) was applied to clusters that displayed thickness values that differed significantly (p < 0.05) from zero. Partial correlations between EDE-Q score and peak thickness values of each cluster were performed in R using the ‘ppcor’ package [Kim, 2015] with age, BMI and WM surface area included as covariates.

### 2.4 Global structural connectivity

To maintain the same resolution as the CT estimates, structural connectivity (SC) was computed at the level of each vertex [Lerch et al., 2006; Raznahan et al., 2011]. Vertex-wise measurements of CT were extracted from each hemisphere across all subjects (N = 163842 vertices per hemisphere), after registration to the fsaverage surface template. Then the CT estimate for each vertex across subjects was correlated to the mean whole-brain CT estimate across subjects. Whereas structural connectivity using CT is typically generated by calculating the pairwise correlations of regional CT cross subjects (i.e. a SC matrix), this estimation of SC approximates the average connectivity of a given vertex. This SC metric has been shown to be highly similar to the average or sum of the connections of a given region or vertex of traditional SC networks [Lerch et al., 2006], and it can be regarded as a summary statistic of how connected a vertex is to all other vertices. Thus, a higher estimate (i.e., a value closer to 1) indicates that, across subjects, CT of a given vertex is generally similar to CT of many other vertices.

### 2.5 Measures of Individual Contribution

As SC is measured across the entire cohort, we estimated an individual’s contribution to the SC by performing a leave-one-out procedure [Saggar et al., 2015]. However, unlike Saggar et al. [2015], who estimated individual contribution by comparing SC matrices using a single Mantel statistic, our vertex-wise estimates of an individual’s contribution were calculated as the difference between the empirical vertex estimate of SC and the vertex estimate of SC without including the CT estimates for a given subject. We call these estimates of individual contribution a leave-one-out (LOO) score. We examined the relationship between the LOO scores and EDE-Q by calculating the Pearson correlation between the LOO score for each vertex and total EDE-Q score. To correct for multiple comparisons, we used bootstrapping (without replacement, N=1000 permutations) to scramble the LOO scores within each subject before calculating the correlations with EDE-Q. Our results are reported using a bootstrapped threshold of p = 0.05.

### 2.6 Exclusions

Four participants were excluded from both global and vertex-wise analyses. Two of these subjects were classified as outliers based on age (values three standard deviations above the mean) and were excluded to eliminate confounding effects of aging on the EDE-Q-CT relationship. The remaining two subjects were excluded because of a technical problem (e.g., corrupted data file), leaving 33 participants for analysis of local CT. One subject was classified as an outlier based on global mean CT (value three standard deviations below the mean), and she was excluded from both global CT and structural connectivity analyses, which both incorporate measurements of average thickness.

## 3. Results

### 3.2 Participants

Within the sample, 12 participants met DSM-IV diagnostic criteria for BN, and five participants met criteria for EDNOS with bulimic symptoms. Inclusion criteria for these groups included: (1) 18.5 ≤ BMI < 29.9 kg/m^2^; (2) regular menstruation for the past six months and (3) at least two objective binge and compensatory episodes in the past 28 days. Remaining participants endorsed either subclinical BN symptoms (e.g., binge eating, driven exercise) or no disordered eating symptoms. No subjects met diagnostic criteria for binge-eating disorder or anorexia nervosa. Eighteen participants reported lifetime treatment for an eating disorder, and two participants used antidepressant medication at the time of the study. Two participants with BN were excluded from analyses for reasons described in Section 2.6.

Previous studies have reported a high level of agreement between the EDE-Q and the EDE when assessing cognitive features of disordered eating in BN [Carter et al., 2001] and BED [Wilfley et al., 1997] patients. Our data support the previously established concurrent validity of the EDE-Q in BN groups. Mean EDE-Q scores were greater in women with clinically significant BN pathology (n = 15; M = 3.65; Range = 1.69 − 5.04) than those without significant pathology (n = 18; M = 1.25; Range = 0 − 2.82). Moreover, EDE-Q score reliably predicted participants with clinically significant BN symptoms versus healthy women (t = 7.11, df = 27.76, bootstrap p < .001). Bootstrapping without replacement (1000 permutations) was performed to assess the robustness of EDE-Q score differences between the two groups.

### 3.3 Association between EDE-Q Score & Global Cortical Thickness

Linear regression analysis indicated a significant negative association between EDE-Q score and global mean CT (β = −0.024, SE = 0.009, *t*(28) = −2.65, p = .013). The effect of age on mean thickness also reached nominal statistical significance (β = −0.009, SE = 0.004, *t*(28) = −2.05, p = .05). The model accounted for 18% of the variability in global thickness (F(3, 28) = 3.32, p = .03, R^2^_*Adjusted*_ = 0.18).

### 3.4 Association between EDE-Q Score & Local Cortical Thickness

Vertex-wise analysis identified a significant, negative association between BN symptom severity and CT in clusters spanning bilateral inferior frontal, posterior temporal and inferior parietal cortices (see Figure 1A & B; Table 2). In the left hemisphere, EDE-Q score was negatively related to CT in the caudal middle frontal gyrus (MFG; r = −0.52, p = .003), medial orbitofrontal cortex (OFC; r = −0.59, p < .001), middle temporal cortex (r = −0.68, p < .001) and inferior parietal lobule (r = −0.51, p = .004). Similar findings were observed in the right hemisphere. Increasing symptom severity was related to cortical thinning in the right superior frontal gyrus (SFG; r = −0.52, p = .003), lateral OFC (r = −.59, p < .001), inferior temporal cortex (r = −.71, p < .001) and a fourth cluster spanning the postcentral gyrus and superior and inferior parietal lobules (r = −0.61, p < .001).

**Figure 1.**
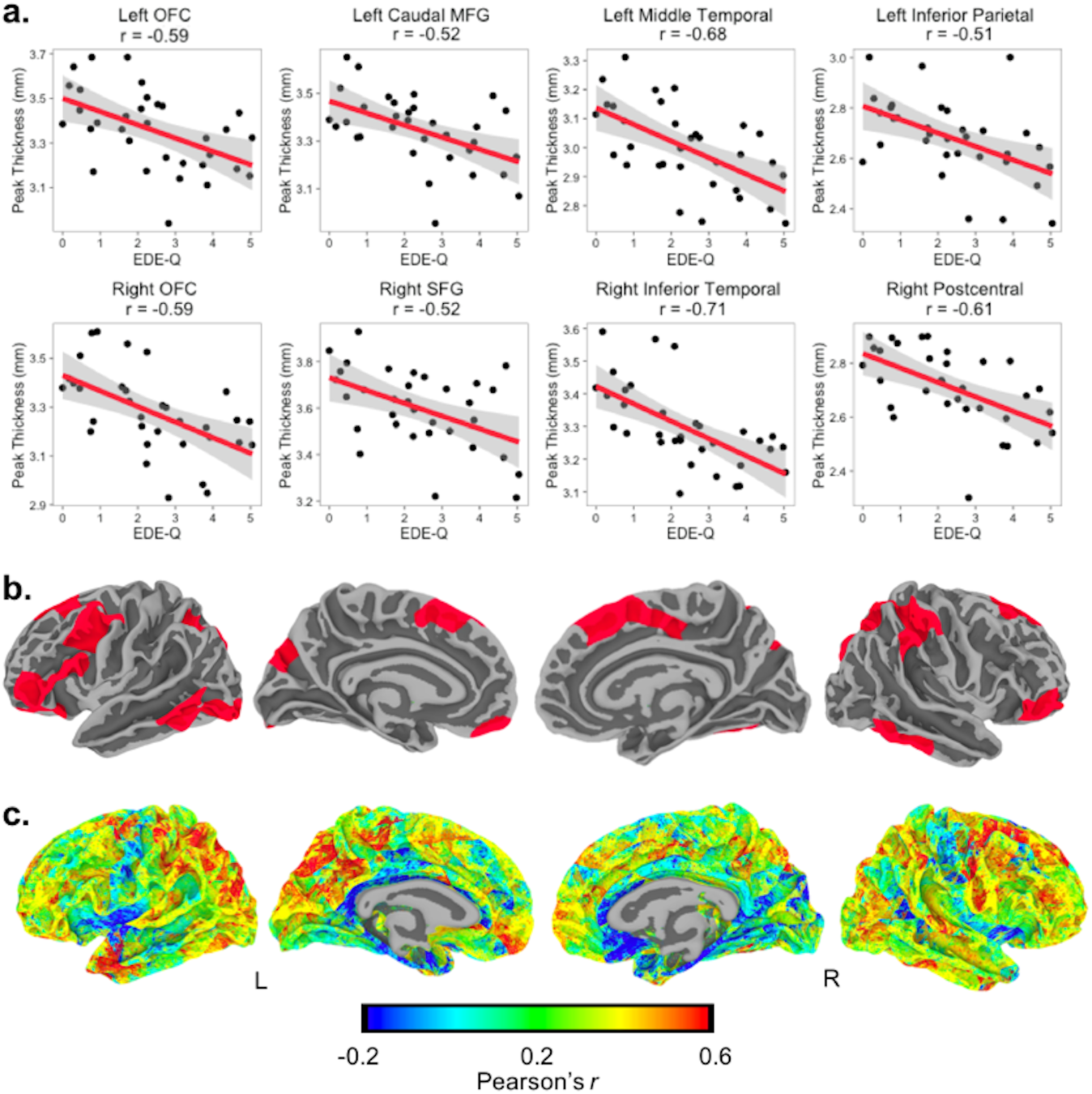
EDE-Q related cortical thinning preferentially occurs in regions with greater global structural connectivity. **a)** Scatterplots depicting partial correlations between bulimia nervosa (BN) symptom severity (quantified as EDE-Q score) and peak cortical thickness (CT) in eight significant clusters. **b)** Surface maps of CT clusters in which increased BN symptom severity was related to greater cortical thinning. Maps were corrected for multiple comparisons at a threshold of p < .05 **c)** Global structural connectivity map of CT, where Pearson’s *r* is the correlation between vertex-wise CT values and mean global CT. Cortical thinning related to BN symptom severity preferentially occurred across vertices with increased global structural connectivity. **Notes:** OFC = orbitofrontal cortex; MFG = middle frontal gyrus, SFG = superior frontal gyrus; Within Figure 1A, Left and Right OFC plots represent clusters following Monte Carlo Null Z multiple comparisons correction at a threshold of p < 0.01, and all remaining plots illustrate corrected clusters at a threshold of p < 0.05.

**Table 2.** Increased bulimia nervosa symptom severity is related to reduced thickness in bilateral frontal, temporal and parietal clusters.

| FreeSurfer Region | Side | Cluster Size | | Peak MNI Coordinates | | | Peak Score | $P_{corr}$ |
| --- | --- | --- | --- | --- | --- | --- | --- | --- |
|  |  | Vertices | Size (mm<sup>2</sup>) | X | Y | Z | t |  |
| Middle temporal | L | 4375 | 2848.70 | -59.5 | -45.4 | -8.8 | -4.13 | 0.003* |
| Caudal middle frontal | L | 5931 | 3402.60 | -33.1 | 10.7 | 28.5 | -3.51 | < 0.001* |
| Medial orbitofrontal | L | 5722 | 3348.97 | -5.3 | 50.1 | -22.1 | -2.64 | < 0.001* |
| Pars orbitalis | L | 1143 | 718.28 | -36.3 | 35.3 | -9.5 | -2.30 | 0.039** |
| Inferior parietal | L | 5247 | 2797.11 | -32.9 | -69.7 | 44.7 | -2.12 | 0.003* |
| Rostral middle frontal | R | 1243 | 850.63 | 36.2 | 54.0 | -6.2 | -3.39 | 0.020** |
| Postcentral | R | 8235 | 3814.20 | 27.2 | -36.0 | 53.2 | -3.35 | < 0.001* |
| Inferior temporal | R | 3221 | 1916.37 | 52.9 | -36.0 | -20.1 | -3.29 | 0.046* |
| Superior frontal | R | 3676 | 1828.49 | 11.4 | -0.9 | 49.8 | -2.96 | 0.044* |

**Notes:** \* = p-value < .05; \*\* = p-value < .01 for Monte Carlo Null-Z simulations for multiple comparisons correction.

### 3.5 Global Structural Connectivity

Estimations of global connectivity of CT at each vertex indicated high SC (i.e., values closer to 1) in cortical regions in the frontal, temporal and parietal lobes. These regions displayed considerable overlap with clusters in which CT was negatively related to BN symptom severity. Moreover, global SC values within these clusters was significantly greater than those contained in 10,000 randomly generated sets of clusters, *t*(57,309) = 74.52, bootstrap p < .0001 (Figure 1C). These results suggest that the cortical thinning associated with increased EDE-Q score preferentially occurs in regions with greatest global SC.

### 3.6 Individual contributions to global CT connectivity scale with EDE-Q score

Using a leave-one-out procedure, vertex-wise individual contributions to global SC (LOO scores) were calculated by taking the difference between the empirical global SC estimates and the global SC estimates generated without a given subject. The distribution of LOO scores was zero-centered such that positive and negative values indicated a high or low contribution, respectively (Figure 2A). The relationship between LOO score and EDE-Q score was spatially distributed across the cortex (Figure 2A), where strong negative correlations (i.e., decreased connectivity in subjects with high EDE-Q) were found in the left lateral OFC, precentral gyrus and inferior temporal cortex (Figure 2B), as well as the right lateral OFC, rostral MFG, inferior temporal cortex and superior parietal lobule. We observed strong positive correlations (i.e., increased connectivity in subjects with high EDE-Q) in the left rostral and caudal MFG and superior parietal cortex and the right SFG, precuneus, superior parietal cortex and fusiform gyrus (Figure 2C). Moreover, approximately 12% of vertices in which EDE-Q was significantly related to LOO score were located within clusters in which EDE-Q was negatively related to CT (as shown in Figure 1B).

**Figure 2.**
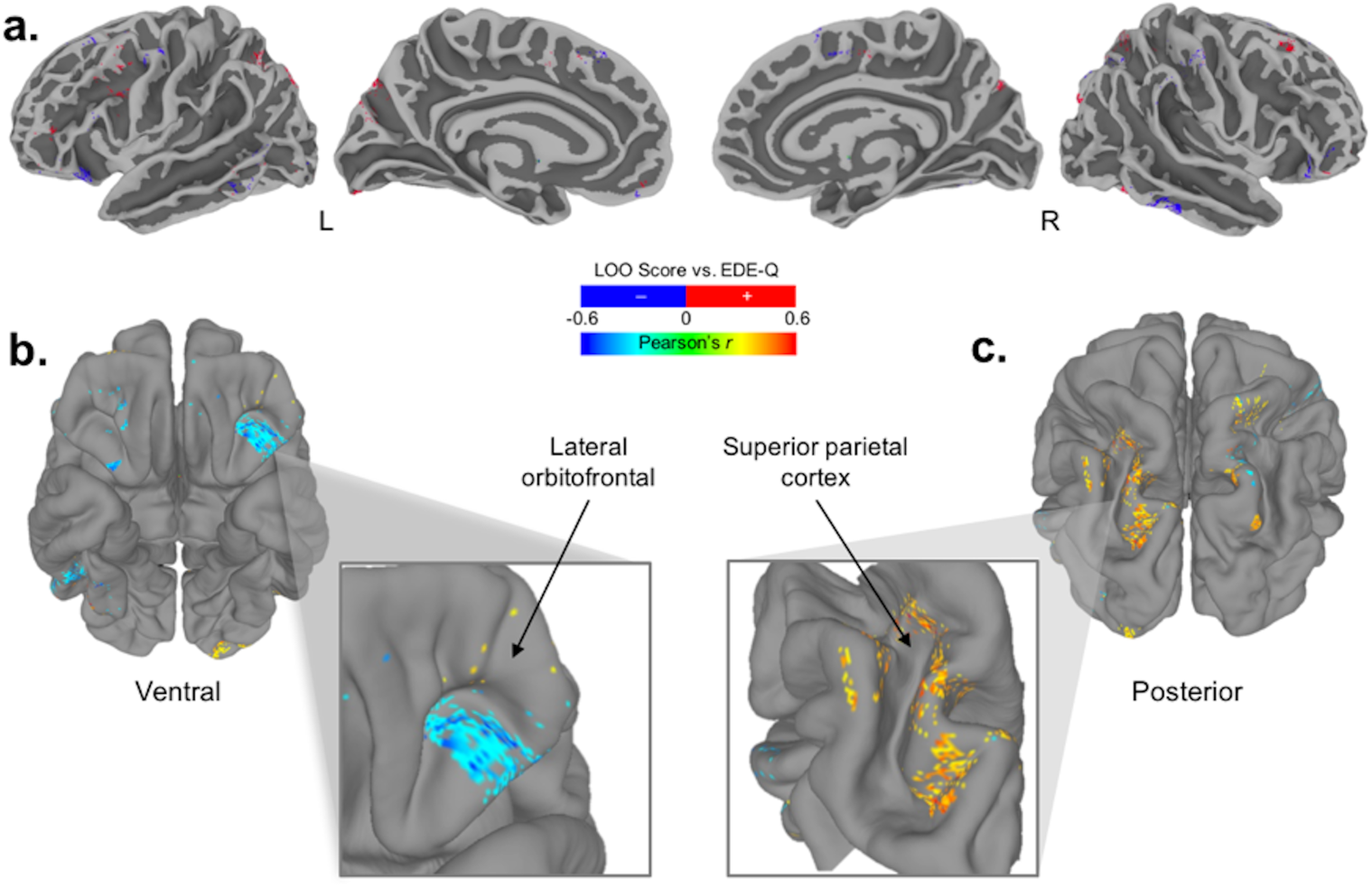
Individual contributions to global structural connectivity scale with EDE-Q score. **a)** Individual contributions to global structural connectivity (LOO scores) plotted on the pial surface. LOO scores were zero-centered; red clusters represent positive contributions to global structural connectivity and blue clusters negative contributions. **b)** LOO score was negatively related to EDE-Q in several regions, including the left OFC. **c)** LOO score was positively related to EDE-Q in vertices spanning the left superior parietal lobule. Correlations between LOO score and EDE-Q (Pearson’s r) were corrected for multiple comparisons using bootstrapping (1,000 permutations; p < 0.05).

## 4. Discussion

We sought to examine associations between BN symptom severity, cortical thickness and accompanying structural connectivity in a sample of adult women who reported a range of BN symptoms. Bulimic symptom severity was quantified using a validated, self-report measure of cognitive and attitudinal aspects of disordered eating, the EDE-Q. Increased EDE-Q score was associated with reduced cortical thickness, both globally and locally in regions spanning the left MFG, right SFG and bilateral orbitofrontal, temporal and parietal cortices. These local associations overlapped considerably with vertices that demonstrated high global structural connectivity of cortical thickness, suggesting that cortical thinning associated with BN symptom severity preferentially occurs in regions with co-varying cortical thickness. Moreover, estimates of each subject’s contribution to global structural connectivity at the group level scaled with EDE-Q score, where both increases and decreases in structural connectivity were related to greater BN symptom severity in a region-dependent manner. Our findings provide novel insight into the anatomical substrates of BN, where both cortical thickness and individual measures of structural connectivity were dimensionally related to symptom severity.

A growing body of literature supports cortical thickness as a robust anatomical feature that is related to underlying cytoarchitecture [e.g., Eickhoff et al., 2005]. Cortical thickness encapsulates the laminar structure of the cortex, and thickness gradients have been shown to complement the brain’s functional sensory hierarchies [Wagstyl et al., 2015]. Although cellular processes cannot be inferred from MRI-derived cortical thickness alone, histological data suggest that decreased dendritic processes [Nakamura et al., 1985] and cell body size [Terry et al., 1987] and increased axonal caliber [Paus, 2010] contribute to gross cortical thinning. To our knowledge, histological brain changes in BN have not been assessed, but a single case study of AN reported increased slender-shaped pyramidal neurons, fewer basal dendritic trunk and spines and reduced cell body size of right precentral gyrus neurons [Neumärker, 1997]. While the precise association between these morphological changes and eating-disordered symptoms is unclear, similar alterations have been associated with brain function in individuals with intellectual disabilities [Huttenlocher, 1991]. This suggests that morphological alterations in the cortex may impact broad cognitive ability. Indeed, cortical thickness has been associated with more specific cognitive attributes, including executive function [Burzynska et al., 2012], attention [Westlye et al., 2011] and impulsivity [Shaw et al., 2011]. Thus, we argue that cortical thickness captures cytroarchitectonic properties of the cortex that are associated with cognition, and assessment of this feature may offer more nuanced insight into the neurobiological bases of BN symptoms.

In eating disorder populations, alterations in cortical thickness may be impacted by the physiological consequences of significant perturbations in feeding and interoceptive signaling. Alterations in the internal milieu are thought to contribute to cortical ‘pseudoatrophy’ in AN, potentially by way of altered hormonal signaling and subsequent glial and neuronal restructuring [Ehrlich et al., 2008; Mainz et al., 2012]. Findings of rapid recovery of cortical thickness in weight-restoring AN [Bernardoni et al., 2016] further substantiate claims that metabolic factors (i.e., alterations in fat mass), as opposed to hydration status or neuronal apoptosis, play a central role in grey matter integrity in AN. However, the extent to which hormonal and metabolic factors contribute to altered cortical morphometry in BN remains unknown, and the relative contribution likely differs from that in AN, where distinct metabolic alterations arise, in part, from low BMI [Misra and Klibanski, 2014] that is not typically observed in BN [Fairburn and Cooper, 1982]. We found EDE-Q-related cortical thinning when controlling for the effects of BMI, so body weight did not appear to have a robust effect on this relationship. It remains possible that biological consequences of chronic binge eating and purging indirectly contributed to the association between BN symptom-severity and global and local cortical thinning. However, our dimensional approach included women with a range of BN symptoms, and only a minority of these women reported self-induced vomiting (n = 5; 15%). We hesitate to draw strong conclusions regarding the role of physiological alterations in the observed global cortical thinning, since local cortical thinning of the eight large clusters may have contributed to the global effect. We would instead cautiously suggest that distorted thoughts about one’s weight and body image in the context of BN are related to cortical thinning, even in the absence of severe disturbances in feeding. This hypothesis aligns with evidence of temporoparietal thinning in patients with body dysmorphic disorder (BDD) relative to healthy individuals [Grace et al., 2017]. Given that BDD is characterized by preoccupation with perceived physical defects, which may include dissatisfaction with weight and body fat (DSM-V, 2013), these findings lend preliminary support to associations between distorted body image and cortical thinning in individuals who do not engage in binge eating and purging.

We observed reduced cortical thickness with elevated BN symptom severity in several regions that exhibited high global structural connectivity of cortical thickness. This finding aligns with previous evidence of co-varying grey matter volume, cortical thickness and surface area of distributed brain regions, a phenomenon termed *structural co-variance* [Alexander-Bloch et al., 2013b; Lerch et al., 2006; Mechelli et al., 2005]. More specifically, alterations in patterns of co-varying cortical thickness have been reported across a range of psychiatric disorders [for review, see Alexander-Bloch et al., 2013a], and our results suggest that coordinated grey matter alterations are also present in women with varying BN symptoms. We assessed structural connectivity using a methodology that has been shown to closely approximate traditional measures of structural covariance [Lerch et al., 2006]. Since structural covariance networks partially recapitulate known functional networks in healthy individuals [Alexander-Bloch et al., 2013b], we aim to interpret the potential relevance of cortical thinning to increased BN symptoms by examining overlap with known functional networks.

Several clusters in which cortical thickness significantly varied with EDE-Q score, notably the MFG, SFG, lateral OFC and inferior parietal lobule, have been identified as components of the frontoparietal control network [Spreng et al., 2010]. A functional intermediary between the dorsal attention and hippocampal-cortical memory networks, the frontoparietal control network is suggested to have a role in integrating information from the external environment with that stored as internal representations [Miller, 2000]. As such, the network contributes to controlled monitoring of information and adaptive task control [Cole et al., 2013; Dosenbach et al., 2007]. Critically, Cole et al. [2014] propose that the network regulates of a variety of brain systems by using feedback control to optimize behavioral outcomes, and impairments in frontoparietal control contribute to a wide range of psychiatric disorders. The authors posit that, in mental illness, the majority of network capacity is devoted to regulating symptoms of the illness, and this reduces the network’s capacity to regulate other domains, such as problem solving in daily life [Cole et al., 2014]. In the context of BN symptom severity, an individual may attempt to regulate negative thoughts about weight and shape by suppressing these. It may be that reduced structural integrity of the network impairs an individual’s ability to discover and implement more adaptive control strategies (e.g., cognitive restructuring). Indeed, prospective studies of women with BN indicate the persistence of distorted thoughts about one’s weight and shape in the absence of behavioral symptoms [Bardone-Cone et al., 2010; Cogley and Keel, 2003], suggesting an impairment in the incorporation of alternative cognitive control strategies. As adaptive cognitive control was not assessed in our study, it would be advantageous for future work to examine associations between cortical thickness, functional connectivity and cognitive control in those who report BN symptoms.

In addition to regions within the frontoparietal control network, we observed cortical thinning of the OFC and the middle and inferior temporal cortex with increased EDE-Q score. The role of the OFC in feeding behavior has been well described [Kringelbach and Radcliffe, 2005; Rolls and Baylis, 1994], and some fMRI evidence suggests hyperactivity of the medial OFC in BN groups when viewing palatable food cues [Uher et al., 2004]. We found reduced cortical thickness in clusters spanning the left lateral and medial OFC and the right lateral OFC. Given that both the medial and lateral OFC are thought to be necessary for the integration of somatosensory information in decision-making processes [Kringelbach and Radcliffe, 2005], it is perhaps unsurprising that cortical thinning of both sub-regions was related to greater BN symptom severity. For example, reduced OFC activity in response to unexpected taste reward has been observed in BN relative to healthy groups during learning [Frank et al., 2011], and recent findings suggest distinct functional connectivity of the anterior cingulate cortex and medial OFC in BN [Frank et al., 2016]. Examination of corticocortical connectivity of the OFC in non-human primates has identified strong reciprocal connections with the temporal cortex [Cavada et al., 2000]. It should be noted that we observed thinning of the left middle temporal cortex, which extended caudally to the lateral occipital cortex, a region that sub-serves visual processing of the human body [Downing et al., 2001]. One possible interpretation may be that, whereas dysfunction of the frontoparietal control network has a central role in mental illness, cortical thinning in these regions relates to cognitions that are relatively specific to BN (e.g., altered taste processing, body image perception).

Our findings represent an important extension of previous case-control studies of brain structure in BN, as our dimensional approach presents the first evidence of cortical thickness alterations in women with a broad range of disordered eating symptoms. Although dimensional models of BN are still debated [see Williamson et al., 2005], some argue for a model where binge eating represents a dichotomous component (i.e., an individual either binges or does not), and body image concerns and drive for thinness exist on a continuum [Williamson et al., 2002]. Thus, our use of EDE-Q score, as opposed to frequency counts of bingeing and purging episodes, may capture aspects of BN pathology that exist at varying degrees in the population. However, while correlations between EDE-Q score and cortical thickness accounted for individual differences, our measurement of global structural connectivity could not be directly related to individual EDE-Q scores. Since quantification of individual differences in cortical connectivity will be critical to relating clinical data to underlying brain structure [Saggar et al., 2015], we used a leave-one-out (LOO) approach to extract vertex-wise, individual contributions to group-level structural connectivity (illustrated in Figure 2A). Correlation of each subject’s EDE-Q score and LOO score at each vertex indicated that, in some areas like the left superior parietal lobule, greater contribution to global structural connectivity was related to increased EDE-Q score (Figure 2C). In other clusters of vertices, high EDE-Q score was associated with less contribution to global structural connectivity (Figure 2B). This can be conceptualized as weaker correlations between cortical thickness in these clusters and global cortical thickness in individuals who report elevated BN symptoms. Negative associations between EDE-Q and LOO scores were clustered in the left lateral OFC and inferior temporal cortex. These findings not only support the notion that individual differences from group-level structural connectivity could represent a ‘biomarker’ for altered brain connectivity [Saggar et al., 2015], but also suggest that both increases and decreases in structural connectivity of cortical thickness are related to BN symptoms, depending on cortical region.

Despite these novel contributions, there were several limitations to the study that may be improved upon in future work. We examined a modest sample size, and assessment of cortical thickness and connectivity within a larger sample of women with variable BN symptoms would better determine the robustness of our findings. In addition, while substance use and psychosis symptoms were exclusion criteria for this study, a more detailed assessment of psychiatric comorbidities (or inclusion of a positive psychiatric control group) would enable examination of anatomical substrates that are specific to disordered eating. More detailed measurement of age of onset, duration and metabolic correlates of BN symptoms would also aide interpretation of possible mechanisms of cortical thickness alterations in women with ED symptoms. Finally, longitudinal assessment of cortical thickness alterations in women with BN symptoms will be critical to advancing knowledge of the role of neurodevelopmental and physiological factors in disease progression.

In summary, our findings of reduced cortical thickness with elevated BN symptom severity preferentially occurred in brain regions with high global structural connectivity, and these regions have been implicated in cognitive processes relevant to BN. We demonstrated that the combination of cortical thickness and structural connectivity measures reflects individual differences in BN symptom severity, which further supports the use of dimensional approaches to determine more nuanced insight into underlying biological mechanisms [Cuthbert, 2014]. Importantly, our findings of bilateral inferior frontal thinning align with those of Marsh et al. [2015] who reported reduced grey matter volume of inferior, middle frontal and central gyri in adolescent and adult women with BN. Inferior parietal lobule thinning in our study also accords with reports of reduced grey matter volume of the region with increased drive for thinness in women with AN and BN [Joos et al., 2010]. Taken together, our results provide novel insight into the neurobiology of bulimia nervosa symptoms, underscoring the importance of a dimensional approach in examination of this debilitating condition.

## Acknowledgements

We would like to thank the participants for their time and commitment to this study. The authors declare that there is no conflict of interest.

